# Fictive Learning in Model-based Reinforcement Learning by Generalized Reward Prediction Errors

**DOI:** 10.1101/2025.06.12.659433

**Authors:** Jianning Chen, Masakazu Taira, Kenji Doya

## Abstract

Reinforcement learning (RL) is a normative computational framework to account for reward-based learning. However, widely used RL models, including Q-learning and its variants, fail to capture some key behavioral phenomena observed in animal experiments, particularly the dynamic switch between model-free and model-based control in the two-step task. A fundamental discrepancy is that learning is restricted to experienced outcomes in the models, whereas biological agents may generalize learning to unvisited options based on internal world models, so-called fictive learning. We propose a simple, brain-inspired fictive learning rule and conduct the rodent two-step task to examine whether fictive learning could explain the observed behavior. The learning rule uses a generalized reward prediction error to update both experienced and non-encountered states and actions. The factual prediction error is scaled by the event correlation inferred from the internal model for fictive update. The generalized reward prediction error might be supported by brain-wide dopaminergic broadcasting. Through simulations, we show that this model reproduces key behavioral traits in the two-step task, including stay probabilities and regression analysis, which common RL models fail to explain. Model fitting validates its superior fit over existing alternatives. Furthermore, the model replicates dopaminergic dynamics observed in the same task. This framework bridges normative RL theory and biological learning, offering new insights into adaptive behavior.

## 1 Introduction

Learning from history to improve future decisions is the key to adaptation. Reinforcement learning (RL) [17] is the canonical theory to describe reward-based learning. An RL agent learns to predict the future outcome from experience and takes the difference between the prediction and the actual outcome, the prediction error, to update the prediction. Despite the great success of applying RL to explain the reward-based behavior in animals, RL-based cognitive models cannot explain some particular phenomena. Particularly, the model-based, model-free RL tradeoff is intensively studied, especially in a two-step task. Many attempts have been made for an integrative computational account for it, yet a gap between the theoretical understanding and real behavior remains [2, 3, 6, 14, 18, 9].

One discrepancy between biological learning and those RL-based cognitive models is the capability of fictive learning. Those models only learn about the experienced actions and states, even when experienced and non-encountered events are often related. By contrast, humans and animals often learns by asking, “If I did something different, what outcome would I have gotten?” [4, 8, 15]. Specifically, they learn about non-encountered events by imagining their potential returns based on the information from experience, which requires an adequate world model that informs the correlation between experienced and non-encountered events [4, 15]. Hence, fictive learning likely co-occurs with model-based RL, but it is usually overlooked in the study of model-based, model-free tradeoff. Additionally, payoffs between options are constantly anticorrelated, which encourages fictive learning, in two-step tasks [2, 3] and other decision making tasks [16, 11]. The win-stay-lose-shift strategy implies that animals might learn to use this anticorrelation to guide their decisions.

Fictive prediction error signals were found in the regions that are responsible for factual reward prediction error computation, including striatum [13, 5], anterior cingulate cortex [10], and orbital frontal cortex [1]. In those regions, neurons encoding different actions and states overlap [16, 12] with an unclear computational implication. Factual prediction error might be generalized as fictive prediction error by the inferred event correlation determined by the co-activation (or mutual inhibition) and overlapping of neurons encoding multiple actions and states in those regions.

Here, we implement fictive learning in model-based RL by the generalized reward prediction error and conduct simulation and animal experiments in a two-step task. This study is novel in three aspects. Firstly, the previous studies derive the fictive error as the outcome difference between the chosen and optimal action [13, 5], whereas the optimal action is often unknown, and computing a separate prediction error is expensive. Our model exploits the factual reward prediction error, which has been computed in factual learning. Literature hypothesizes the fictive error as the endogenous supervisor that is independent of the standard RL updating, yet has not verified the model [13, 5]. We propose a simpler and more plausible version with evidence of cognitive modeling. Thirdly, previous studies design the experiment similarly to the stock market, and the anticorrelation between actions, buying and selling, is explicit. In our experiment, animal has to learn such anticorrelation, allowing us to examine whether fictive learning would naturally arise in reward-based learning.

In the following sections, we first describe the experiment design and model. We then simulated existing models without fictive learning to show that they fail to explain the experimental result. Next, we show that a fictive model-based RL fits the observation by simulation, followed by the explanation of action updating. Then, the model fitting confirmed that the fictive MB model fit the behavior better than others. Finally, we conclude with a discussion of the implications and hypotheses for further validation of the model.

## 2 Methodology

### 2.1 Experiment design

We trained 10 mice (C47/BL background) with the two-step task (Fig. 1a) [2, 3]. The mice freely chose between the left and right options in ~ 75 % trials (mice were forced to choose left or right in the rest of the trials), which led to either an up or a down state with either common (80%) or rare (20%) probability. The transition probability matrix was fixed between subjects and counterbalanced across subjects. That is, the left option commonly leads to the up state in some animals, and vice versa in others. Reward is delivered probabilistically at each state, and the reward probabilities of up and down states are changed between 80%/20%, 20%/80%, 50%/50%, termed as up, down, and neutral block (Table. 1). The reward settings changed block-wise. In a non-neutral (up or down) block, block changes after 5 - 15 trials once the exponential moving average of the correct response (i.e., the option that commonly leads to the state with higher reward probability) in the last eight free trials reached 0.75. The reward settings changed after 20 - 30 trials in the neutral block.

**Fig. 1:**
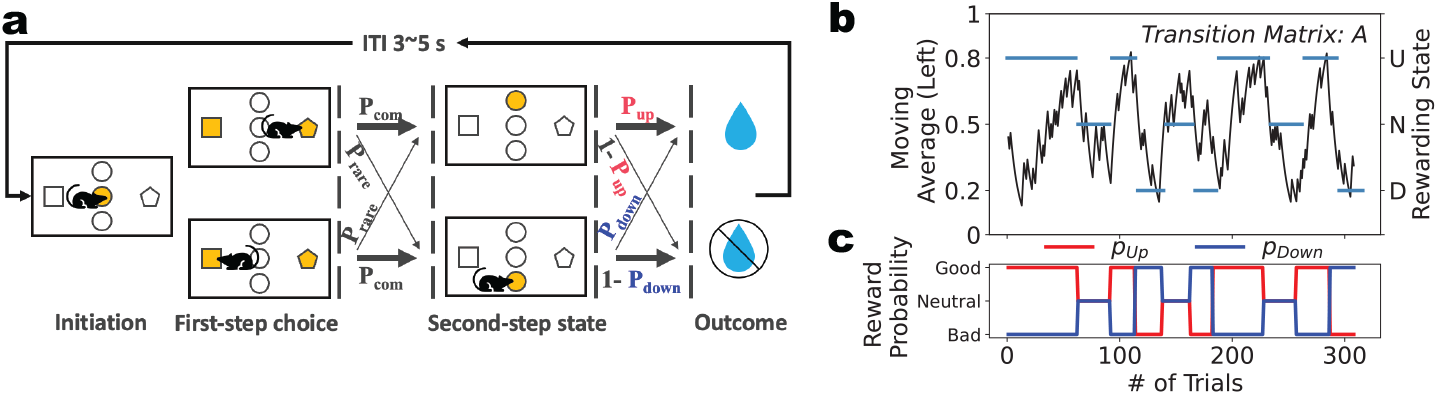
**a**, Task structure. After initiation, a first-step choice between left and right is presented, followed by up or down state with either common (80%) or rare (20%) probability. Two states are rewarded with different reward probabilities. **b**, Example behavior. The exponential moving average of animal choice (black line) traces the reward setting (blue bar). c, reward settings. Reward probabilities change in blocks anticorrelated.

### 2.2 Model description

We included 11 base models, including model-free (MF) and model-based (MB), and mixture models. We then augment those models with fictive learning.

#### Base models

In the task, the agent chooses an action *a* ∈ (*left, right*), which leads to the second-step state *s* ∈ (*up, down*), where the outcome *r* ∈ (0, 1) is delivered. Agents learn the action value *Q*(*a*) differently, yet the action selection follows the softmax function.

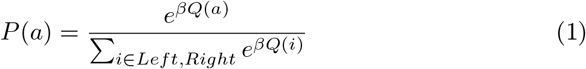

The model-free category includes the model-free model with eligibility trace, MF(lambda), and its two variants, direct model-free, MF, and model-free with memory model, MF(memory). The MF(lambda) agent updates its action value of chosen options *Q*_*mf*_ (*a*) and state value of experienced state *V* (*s*) as follow,

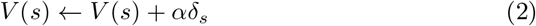

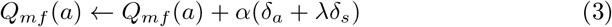

In Equation (2) and (3), *δ*_*a*_ = *V* (*s*) − *Q*_*mf*_ (*a*) and *δ*_*s*_ = *r* − *V* (*s*) are the prediction error for action and state value. The model is termed MF and MF(memory) when the eligibility trace *λ* is 1 or 0, respectively.

The model-based category includes the model-based, MB, and the Bayesian hidden state model, hidden state. Agents exploit the learned model, the transition matrix between actions and state (*P* (*s* | *a*)). The MB agent learns the state value by Equation (2) and then computes the action value by,

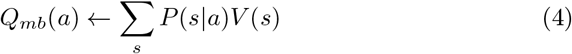

The hidden state agent assume that there are two hidden states in which either of the two second-state states is better and update the beliefs of being one hidden state (*h* ∈ *h*_*up*_, *h*_*down*_) using Bayesian inference [3].

Specifically, the agent estimates the *P* (*h*_*up*_) by,

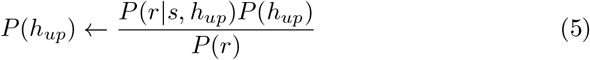

where the likelihood *P* (*r* |*s, h*_*up*_) is the reward probability in experiment (Table. 1).

The marginal likelihood *P* (*r*) is calculated as,

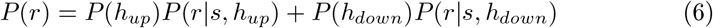

The agent might assume that the environment would change with a certain probability *τ*. Hence, the posterior is updated as,

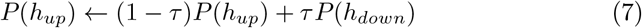

Therefore, the state values are updated as,

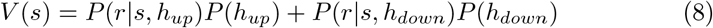

And the action is updated as in the MB model. We include an extra variant for model comparison, the asymmetric hidden state model. In this model, the agent treats the omission in up and down states as the same observation by using the likelihood table (Table 2), while other components remain the same.

**Table 1:** Reward Probability.

| Block | Up block |  | Down block |  |
| --- | --- | --- | --- | --- |
| Outcome | Reward | Omission | Reward | Omission |
| Up state | 0.8 | 0.2 | 0.2 | 0.8 |
| Down state | 0.2 | 0.8 | 0.8 | 0.2 |

**Table 2:** Reward Probability.

| Block | Up block |  | Down block |  |
| --- | --- | --- | --- | --- |
| Outcome | Reward | Omission | Reward | Omission |
| Up state | 0.4 | 0.5 | 0.1 | 0.5 |
| Down state | 0.1 |  | 0.4 |  |

A hybrid model consists of multiple model-free and model-based models, and the action value is computed as,

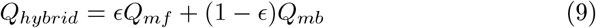

#### Fictive learning

The fictive learning is implemented by updating the state value of the unvisited state (*V* (*s*_*−*_)) and the action value of the unchosen option (*Q*(*a*_*−*_)), by the reward prediction error from visited state (*V* (*s*_+_)) and chosen option (*Q*(*a*_+_)) in Equation (2) and (3). The proportion of updating depends on the inferred event correlation of reward probability between states *η*_*s*_ (*η*_*s*_) and options *η*_*a*_ ((*η*_*a*_)) by,

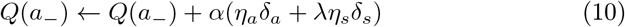

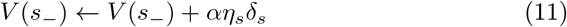

The event correlation factors are zero when the agent believes the reward probability of two actions and states change independently (as in MB model), negative if anticorrelated, and positive if changing in the same direction. In model simulation and fitting, we simplified the model in two aspects. Since the transition matrix was fixed and well-instructed, we assumed that the agent believes the correlations between states and actions are the same (i.e., *η*_*s*_ = *η*_*a*_ = *η*). For the same reason, we assume that the agent already learns the correlation, and *η* is a fixed hyperparameter.

### 2.3 Model Simulation

To examine which model will exhibit behavior similar to the experimental observation, we simulate agents listed in Table 3. To cover a wide range of hyperparameters, we add a noise term *noise ~ 𝒩* (0, 0.05 ×|*hyperparameter* |) for each agent. For each model, we simulated 15 agents for 20 sessions, and each session contained 400 trials, which is the typical length in animal experiments.

**Table 3:**
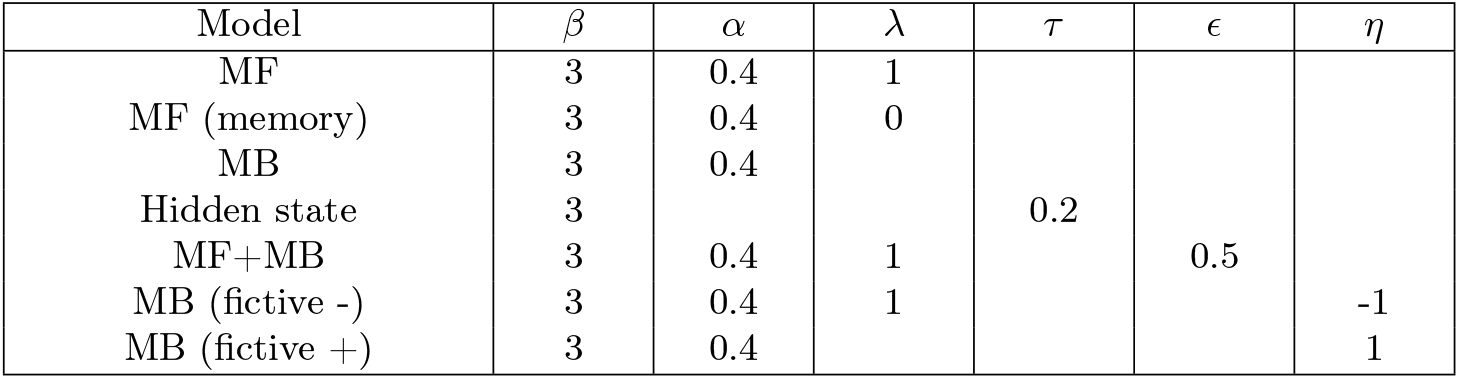
Model and hyperparameter settings.

**Table 4:** Action updating table.

| (a) MB |  |  |  |  | (b) MB (fictive -) |  |  |  |
| --- | --- | --- | --- | --- | --- | --- | --- | --- |
|  | CR | RR | CN | RN | CR | RR | CN | RN |
| $\Delta Q(a_+)$ | $0.8\alpha\delta_s$ | $0.2\alpha\delta_s$ | $0.8\alpha\delta_s$ | $0.2\alpha\delta_s$ | $0.6\alpha\delta_s$ | $-0.6\alpha\delta_s$ | $0.6\alpha\delta_s$ | $-0.6\alpha\delta_s$ |
| $\Delta Q(a_-)$ | $0.2\alpha\delta_s$ | $0.8\alpha\delta_s$ | $0.2\alpha\delta_s$ | $0.8\alpha\delta_s$ | $-0.6\alpha\delta_s$ | $0.6\alpha\delta_s$ | $-0.6\alpha\delta_s$ | $0.6\alpha\delta_s$ |

### 2.4 Model fitting procedure

We implement the Bayesian fitting with Rstan 2.32.6 with 4 MCMC chains for 2000 iterations (500 warm-up runs). We included three MF and two MB models, six hybrid models, an asymmetric hidden state model, and seven fictive learning models. The hidden state model implicitly implements fictive learning by anticorrelation; we did not add fictive learning to those base models in which the hidden state model is involved to prevent confounding. The model fitting was performed for each subject and session to account for high subject/session-level variability. BIC score was used for model evaluation.

### 2.5 Analysis method

Analyses used custom Python, R, and Matlab scripts. For normally distributed data, we use the paired *t*-tests when within-subject comparison with equal sample size and unpaired *t*-tests otherwise. Otherwise, we used Wilcoxon signed-rank tests and Mann-Whitney U-tests, respectively.

#### Generalized linear mixed model

We built the generalized linear mixed model (GLMM) using *fitglme* (Matlab 2023b) to predict the stay/switch behavior in free trials with the logit link function. All included variables were assumed to vary between subjects, and a full random effects matrix with subjects as grouping factors was included for all variables and the intercept. The model structure is,

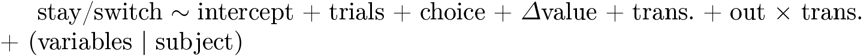

The variables were coded as:

- stay/switch: 1 if the animal stayed at the same choice as the last trial and 0 otherwise.
- trials: number of trials experienced in the session.
- choice: previous action, 0.5 if the previous choice was left, −0.5 otherwise.
- *Δ*Value: inferred value difference. The estimated difference in reward probabilities between the chosen and unchosen options prior to the current trial.

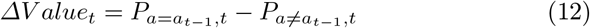

where,

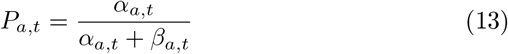

where,

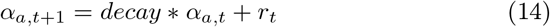

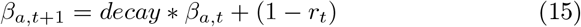

The *decay* is set as 0.5 to ensure the results’ generalizability.
- Trans.: 0.5 if the previous transition is common, −0.5 if rare.
- Out × Trans.: 0.5 if the previous trial was a common/reward transition or a rare/omission transition, −0.5 otherwise.

## 3 Result

### 3.1 Existing model fails to explain the experimental result

10 mice were tested to perform 17.600 *±* 3.720 sessions, and were able to perform 425.500 *±* 57.642 trials and completed 8.290 *±* 1.645 of non-neutral blocks per session. Animals learned to optimize the choice (Fig. 1b) with 64.95% of correct choices.

In simulation, models from the MB and MF classes show distinct stay probabilities and GLMM coefficients (Fig. 2a). MB and hidden state model switch frequently after a rare/reward and common/no-reward trials, leading to the positive coefficient of interaction of outcome and transition in GLMM (Out × Trans.. MB: *β* = 0.684, SE = 0.026, *t* = 25.925, *p* < 0.001; hidden state: *β* = 0.678, SE = 0.016, *t* = 43.616, *p* < 0.001). Such a tendency reverses or disappears in the stay probability and GLMM for MF (*β* = −0.072, SE = 0.017, *t* = −4.183, *p* < 0.001) and MF (memory) (*β* = 0.018, SE = 0.015, *t* = 1.184, *p* = 0.236). The stay behavior depends heavily on the reward prediction of the chosen one over the unchosen ones based on the reward history (i.e., *Δ*Value) in model-free models (MF: *β* = 2.691, SE = 0.026, *t* = 102.09, *p* < 0.001; MF(memory): *β* = 0.920, SE = 0.020, *t* = 45.136, *p* < 0.001) than model-based models (MB: *β* = 0.922, SE = 0.020, *t* = 45.738, *p* < 0.001; hidden state: *β* = 0.226, SE = 0.018, *t* = 12.679, *p* < 0.001), as it essentially captures the direct reinforcement of the outcome on the action. Besides, all models show a similar tendency to repeat actions (i.e., intercept) and no or a marginal effect of transition type (Trans. MF(memory): *β* = 0.095, SE = 0.018, *t* = 5.370, *p* < 0.001).

**Fig. 2:**
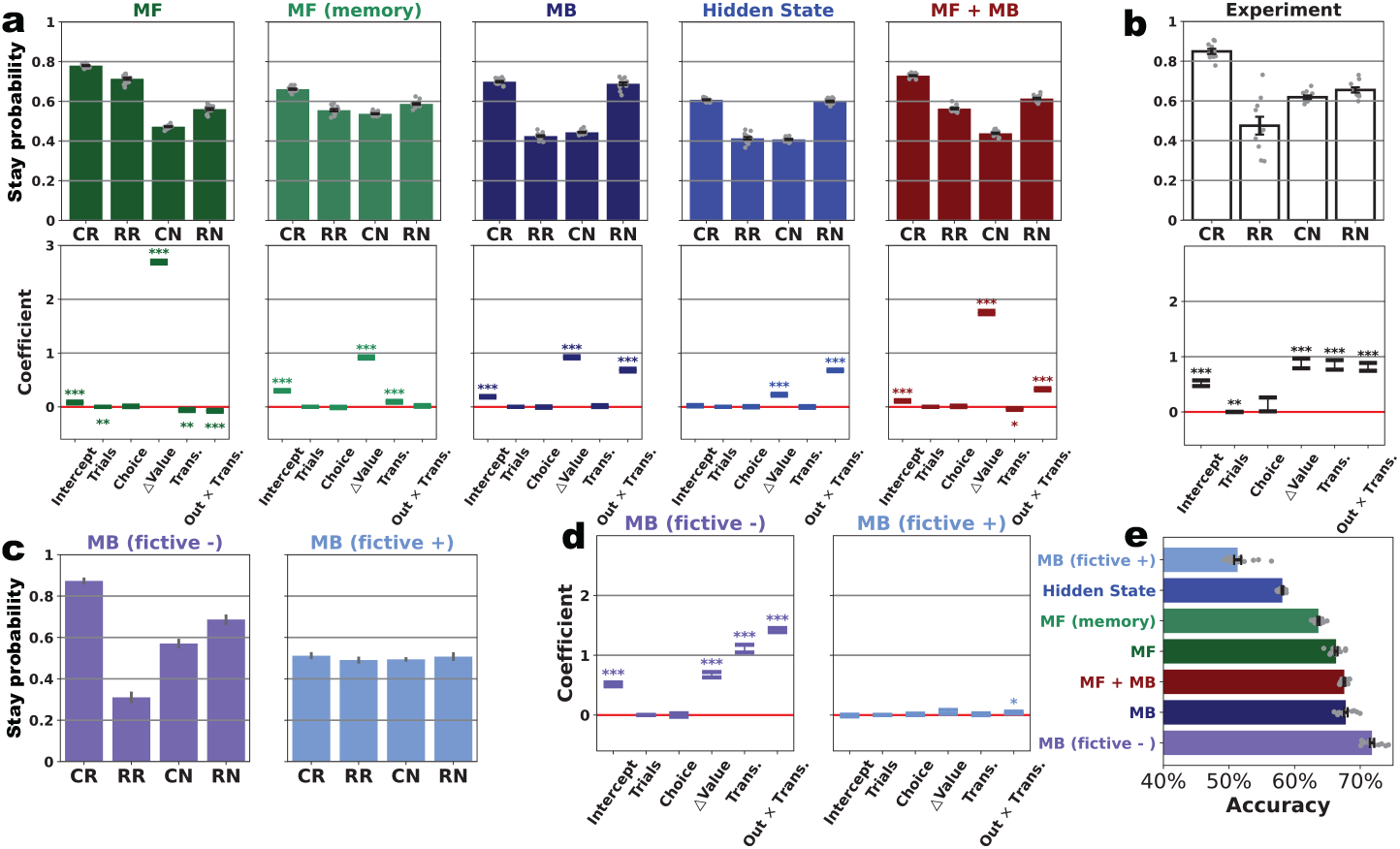
**a, b**, Behavior in simulation (a) and experiment (b). **Top**, the stay probability after trials with different outcome-transition pairs. Dots show each subject, and the error bar shows the between-subject mean *±* s.e.m.. **Bottom**, GLMM result. The error bar shows the estimated coefficient *±* SE, and the star represents the significance. *, 0.01 ≤ *p* ≤ 0.05; **, 0.001 ≤ *p* ≤ 0.01, and *** ≤ 0.001.**c, d**, stay probability (c) and GLMM result (d) of fictive learning agent. **e**, The accuracy of included agents.

Animal behavior is different from the above agents (Fig. 2b). Animals show the highest stay probability after the common/reward trial (84.923 *±* 3.910%), resulting from the effect of value history (*Δ*Value: *β* = 0.623, SE = 0.086, *t* = 7.265, *p* < 0.001). However, animals were likely to switch following the rare/reward trial, and the stay probability is marginally lower after common/no-reward trials (61.863 *±* 2.763%) than rare/no-reward trials (65.513 *±* 3.973%)(*stat* = 5.00, *p* = 0.020, Wilcoxon test), suggesting the effect of the interaction of outcome and transition type (Out *×* Trans.: *β* = 0.613, SE = 0.061, *t* = 10.080, *p* < 0.001) and the involvement of model-based learning. Besides, transition type strongly modulates the stay probability, mainly after a rewarded trial, whereas it only has a subtle effect after the unrewarded trials, leading to a significant positive coefficient of transition type in GLMM (Trans.: *β* = 0.666, SE = 0.062, *t* = 10.696, *p* < 0.001).

The existing models cannot replicate the observation. Animals’ stay probability after common/reward trials is higher than all model predicts. Animals are more likely to stay in rare/reward trials than in common/no-reward trials. By contrast, the model-based models showed the equal stay probability in two cases, and the model-free and hybrid models showed the opposite pattern. Therefore, none of those models shows a strong positive coefficient of transition type.

### 3.2 Fictive learning agent fits the experiment result

Including fictive learning with anticorrelation (i.e., *η* = −1), MB(fictive −) generates behavior similar to animal behavior in the stay probability (Fig. 2c) and the GLMM coefficients (Fig. 2d). This model also achieves the highest accuracy (71.793 *±* 1.252 %)(Fig. 2e). By contrast, MB(fictive +) (i.e., *η* = 1) shows seemingly random behavior, yet GLMM reveals a significant coefficient of outcome transition type interaction (*β* = 0.041, SE = 0.017, *t* = 2.489, *p* = 0.013).

We examine why MB(fictive −) behaves differently from MB by analyzing how action value is updated. As an example, we domesticate the value updating after common/reward trials, assuming the agent chose the left (*L*) and visited the up state (*D*) (Fig. 3a,b). The action value of chosen (*Q*(*a*_+_)) and unchosen options (*Q*(*a*_*−*_)) are updated via the transition matrix in MB and MB(fictive −) by different magnitudes (Table. 4). The MB model updates the *Q*(*a*_+_) with *δ*_*s*_ scaled by 80% common probability and *Q*(*a*_*−*_) by 20% rare probability(Fig. 3c).

**Fig. 3:**
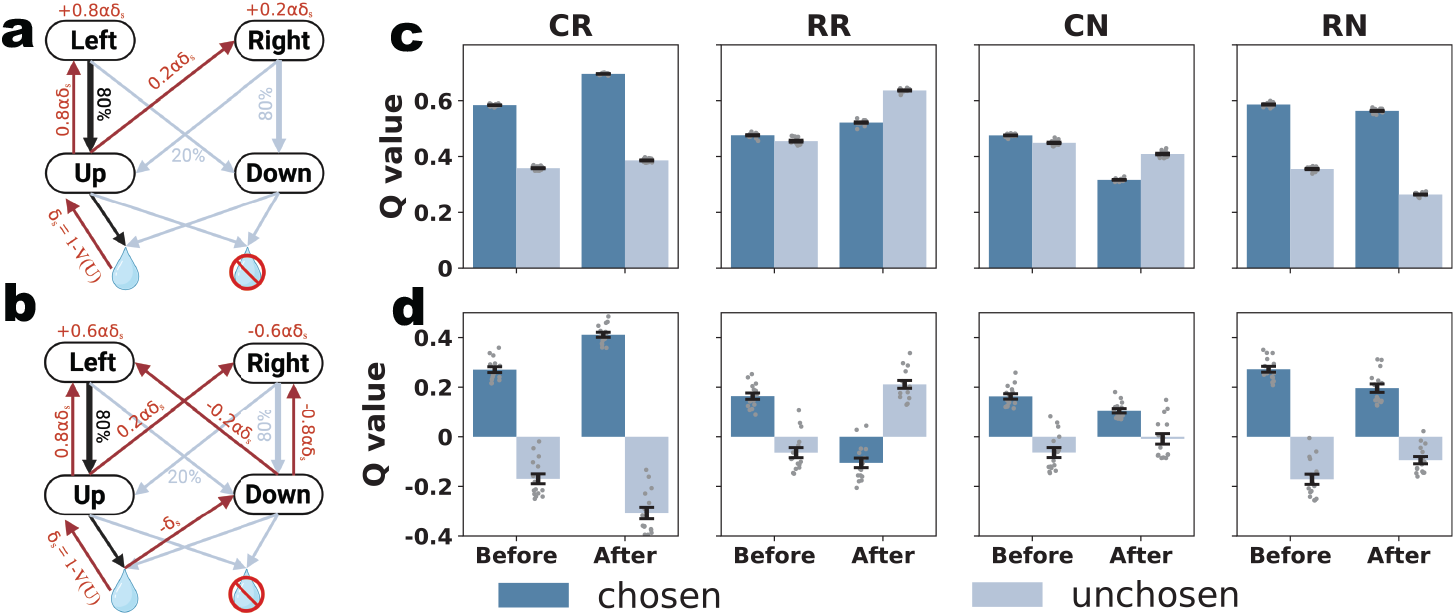
**a,b**, The action value update in common/reward trials. When the left option was chosen, followed by a common transition to the up state, and a reward. Action values of chosen and unchosen options get updated by 0.8*αδ*_*s*_ and 0.2*αδ*_*s*_ in MB (a) and 0.6*αδ*_*s*_ and −0.6*αδ*_*s*_ in MB(fictive-) (b)(Created in BioRender. https://BioRender.com/e8qv5x0). **c**,**d**, Action updating in MB (a), and MB(fictive −) (b) in four transition/outcome pairs.

In MB(fictive −), action value updates by two opposite *δ*_*s*_ via the transition matrix, resulting in the simultaneous reinforcing of *Q*(*a*_+_) and fictive punishing *Q*(*a*_−_)(Fig. 3d).

After a rare/reward trial, preference reversal happens with distinct rationales in the two models. The MB model (Fig. 3c) learns to increase the action value of both options, but increases the *Q*(*a*_*−*_) for large magnitudes, leading to the takeover in action value and a switch choice. By contrast, MB(fictive −) (Fig. 3d) decreases the *Q*(*a*_+_) but increases the *Q*(*a*_*−*_). Thus, the takeover in action value is more substantial in MB (fictive −) and leads to a lower stay probability than MB (Fig. 2a,c).

After common/no-reward trials, two models update the action value by the negative prediction error differently. In the MB model, *Q*(*a*_+_) decreases dramatically, yet *Q*(*a*_*−*_) drops modestly, leading to the preference reversal (Fig. 3c). In MB(fictive −) model (Fig. 3d), *Q*(*a*_+_) decreases and *Q*(*a*_*−*_) increases. Since the agent and animals performed the task well, getting a reward omission after a common transition, an incorrect choice, is not due to insufficient learning, but happened because omission happened with 20% probability, or block change. In either case, *Q*(*a*_+_) should be substantially higher than *Q*(*a*_*−*_), and hence one-shot updating is not enough to trigger the preference reversal but only attenuate the difference between action value.

After rare/no-reward trials, in MB model (Fig. 3c), an omission causes a negative prediction error, leading to a notable decrease in *Q*(*a*_+_) and a slight decrease in *Q*(*a*_*−*_) without preference reversal. However, in the MB(fictive −) model (Fig. 3d), a rare transition usually directs the agent to an infavored state with negative state value by fictive punishment (i.e., given the negative correlation between state, since the visited state gives fascinate outcome, being unvisited state might give a punishment), implying the prediction error can be positive in some cases. Hence, the action value difference was equalized, but no preference reversal happened.

### 3.3 Fictive Model-based agent fits the observation better

The Bayesian modeling and model comparison suggest fictive learning better captures the observed behavior. The comparison between base models shows that the MB model provides better fits in general (*Δ* BIC: 38.587 *±* 28.149) (Fig. 4a,b), suggesting that model-based learning is indeed dominant.

**Fig. 4:**
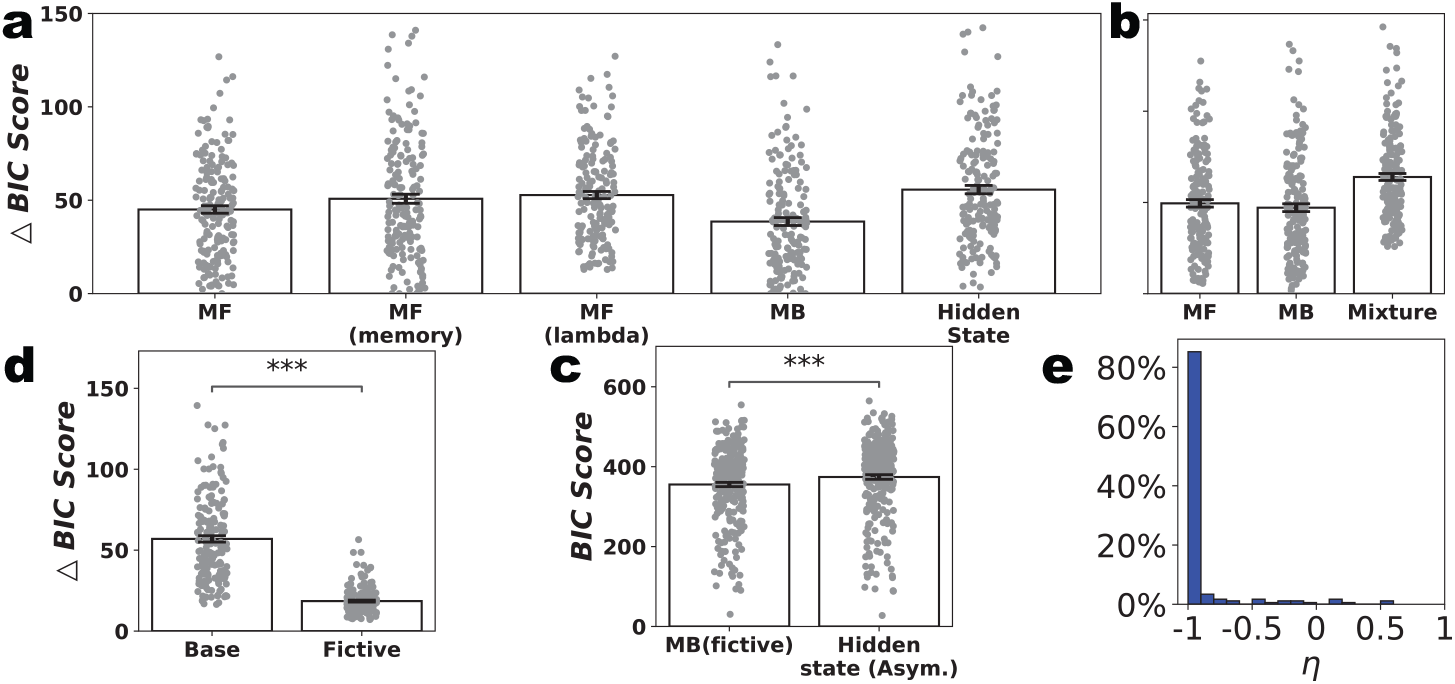
**a**, Model comparison of single base models. The *Δ*BIC score is the BIC score normalized by the BIC score of the winning model per subject/session. The MB model achieves the lowest *Δ*BIC score. **b**, Model comparison of MF, MB, and mixture classes. MB classes fit the data better. **c**, Model comparison between all base models and their variants with extra fictive learning. Extra fictive learning improves model fit. **d**, Model comparison between MB (fictive) and asymmetric hidden state model. MB (fictive) fits the data better in general. **e**, estimated event correlation parameter *η. η* tends to be around −1.

Adding fictive learning causes a significant decrease in BIC score (Fig. 4c) (base model: 56.970 *±* 25.994; fictive agent: 18.545 *±* 8.201; *stat* = 0, *p* < 0.001, Wilcoxon test). A hidden state model that learns differently from reward and omission could also predict a similar behavior pattern [3]. Yet, MB(fictive) shows a better fit than it suggested by absolute BIC score (Fig. 4d) (asymmetric hidden state model: 374.183 *±* 97.681; MB (fictive): 355.407 *±* 90.124; *stat* = 9521.000, *p* < 0.001, Wilcoxon test), despite having the same number of hyperparameters. Consistent with simulation and task setting, estimated *η* is negative (−0.906 *±*0.248) (Fig. 4e), implying that animals learned the event anticorrelation.

### 3.4 Prediction error explains the striatal dopaminergic activity

The prediction error from MB(fictive −) is consistent with the dopaminergic (DA) activity in the nucleus accumbens in mice performing the same task [3]. Strital DA dynamics are believed to signal the prediction error modulated by the last outcome. [3] reported the reversal in the coefficient of the last outcome predicting the DA activity in the current trial. Specifically, it was negative when the second-step state was revealed but positive during the outcome period, if the presented state was the same as in the last trial. Yet, it shows a negative-to-positive reversal when experiencing the state that was not visited before, for which the classical MB model fails to reproduce.

To examine if our model would reproduce this phenomenon, we derive the prediction error as a proxy of the dopaminergic signal and predict it by the last outcome. Transition prediction error is the difference between the state value before and after the updating in the last trial of the current presented state (*V* (*s*_+_, *t*) *− V* (*s*_+_, *t −* 1)), and the outcome prediction error is the difference between the outcome and the state value *r*(*t*) *− V* (*s*_+_, *t*). In Fig. 5a, when experiencing the same state, all models replicate the same reversal observed in real DA activity. However, when experiencing the different state (Fig. 5b), MB(fictive −) reproduces the observed pattern. Furthermore, MB(fictive +) predicts the positive-to-negative reversal in both cases. This analysis suggests that the fictive learning might underlie the behavior, but also the neural computation.

**Fig. 5:**
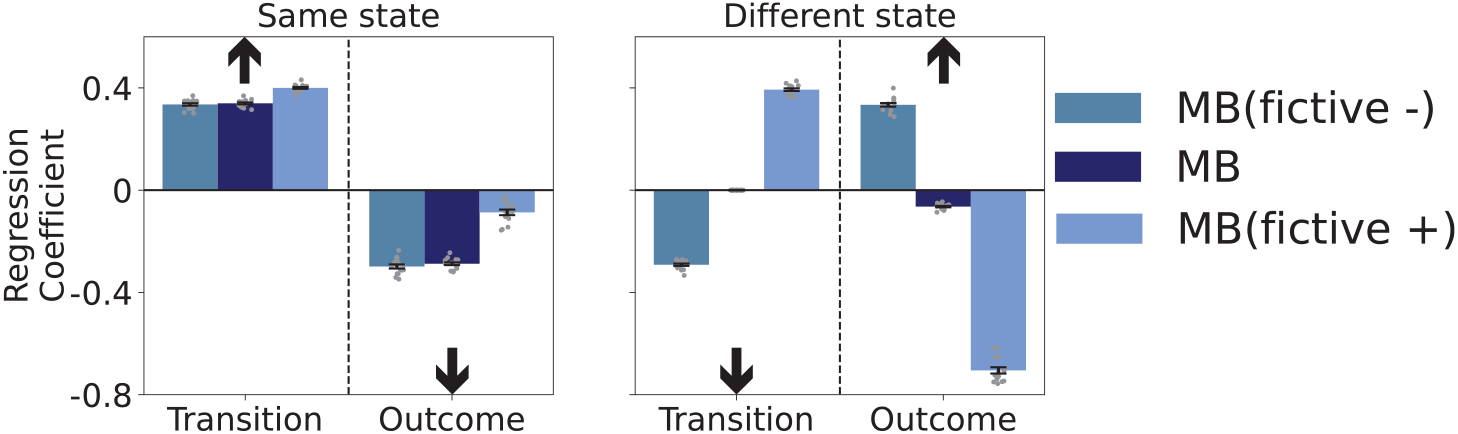
**a, b**, The coefficient of last outcome predicting the hypothetical dopamine signal derived from prediction error in MB (fictive −), MB and MB (fictive +), when the same (a) or different (b) second-step state is presented. All three models predict the same pattern when the same state is presented. However, only MB (fictive −) predicts the same pattern as observed in dopamine release when the different state is presented. Arrows show the direction of the coefficient predicting the dopamine signal at the nucleus accumbens recorded in [3].

## 4 Discussion

This study introduces a novel model that integrates fictive learning with the model-based RL model. This model achieves the best performance against other alternatives in the two-step task. It also reproduced the observed behavior and DA dynamics better. Here, we discuss the merits and limitations of the model, the possible biological mechanisms, a theoretical comparison with competing models, and testable hypotheses for further validation of this model.

### Merits and limitations of fictive model-based model

The fictive model-based model learn the task efficiently. The simultaneous reinforcing and punishing mechanism facilitates learning, leading to the highest accuracy. Fictive learning learns more efficiently from feedback by exploiting the task structure. After understanding the task, the fictive learning agent could stay at the optimal choice more deterministically by enlarging the contrast between choices two-fold compared to other agents, and avoid exploration by explicitly updating the unvisited state and unchosen choice.

A fictive model-based model also fit the observed behavior in the two-step task better. The conventional view believes that animals balance between the model-based and model-free RL to solve the task [6, 2]. The presented model provides a simpler, more integrated, and plausible interpretation. However, it does not imply that no model-free component is involved. In this study, we only included the data after training, where animals are assumed to learn the transition matrix and event correlation. Model-free might dominate when uncertainty elevates [7] during training or changes in event correlation. How event correlation is learned, and whether it drives the switch between model-free and model-based systems, could be a fruitful direction.

### Hypothetical biological mechanism

The fictive model-based model reproduces the prediction error that fits the dynamics of dopamine in the nucleus accumbens [3], suggesting the brain might execute a similar computation. Such fictive learning could naturally arise from Hebbian and anti-Hebbian plasticity. The co-activation or mutual inhibition between neurons that represent experienced and non-encountered events develops from learning and allows for fictive learning. Furthermore, since anticorrelation between options is embedded in most behavioral tasks, the observation that some neurons represent different options or states [16] might be a result of fictive learning.

### Comparison with existing models

Our model performs similarly to an asymmetric hidden state model [3], yet the presented model provides a more generic and realistic computational account. Both our model and the asymmetric hidden state model reproduce the observed behavior and neural activity. However, the significant difference in stay probability after common/no-reward and rare/no-reward trials is not consistent with the assumption that agents treat the reward omission in up and down states as the same observation in the asymmetric hidden state model. Hence, our model provides a lower BIC in model comparison. The hidden state model implicitly implements the update of unvisited states and unchosen options by assuming that one state (and hence option) is better than the other. Yet, it only allows for the negative fictive update, whereas our model provides a more generic rule allowing for negative, positive fictive update, and classical model-based update.

In asymmetric hidden state computation, animals must obtain knowledge of a set of reward probabilities. It is already heavy in a two-step task, and will be overwhelming in a more realistic setting with more options and states. However, our model only requires the prediction error to be broadcast and a rough estimate of event correlation, which is computationally cheap and biologically plausible.

### Testable hypotheses for further validation

Here are testable hypotheses to examine whether the brain implements a fictive model-based RL. In behavior, when correlation changes from negative to positive, our model predicts the stay probability, from the pattern observed here, to have an inverted-U shape from the classical model-based model, to seemingly random. The GLMM result would also change as the coefficient of transition type and value history diminishes. In neural activity, the fictive learning predicts that the coefficient of the last outcome on dopamine release is modulated by whether the same state is presented. This modulation disappears when event correlation is positive. Besides, co-activation between neurons or the proportion of neurons representing multiple options or states gets weaker or smaller when correlation is weakened.

## Conclusion

In conclusion, we integrate model-based RL with fictive learning and conduct in-silico and animal experiments to examine its ability to learn faster and explain animal behavior. We found that fictive learning facilitates learning while being algorithmically simple and biologically plausible. Model simulation and fitting show that it describes the behavior and dopamine dynamics in the two-step task better than the existing model. The presented result contributes to filling the gap between biological learning and the current RL theories.

## Acknowledgments

We acknowledge Thomas Akam and Mark Walton for their helpful advice on setting up the behavioral task. This study was funded by JSPS KAKENHI Grants 16H06563 and 23H04975, and Research Support of Okinawa Institute of Science and Technology Graduate University to K. D.

